# Adaptive rewiring based on diffusion balances stability and plasticity in weighted networks while evolving ‘brain-like’ structure

**DOI:** 10.1101/723353

**Authors:** Ilias Rentzeperis, Cees van Leeuwen

## Abstract

Activity dependent plasticity is the brain’s mechanism for reshaping neural connections. Representing activity by graph diffusion, we model plasticity as adaptive rewiring. The rewiring involves adding shortcut connections where diffusion on the graph is intensive while pruning underused ones. This process robustly steers initially random networks to high-levels of structural complexity reflecting the global characteristics of brain anatomy: modular or centralized small world topologies, depending on overall diffusion rate. We extend this result, known from binary networks, to weighted ones in order to evaluate the flexibility of their evolved states. Both with normally- and lognormally-distributed weights, networks evolve modular or centralized topologies depending on a single control parameter, the diffusion rate, representing a global homeostatic or normalizing regulation mechanism. Once settled, normally weighted networks lock into their topologies, whereas lognormal ones allow flexible switching between them, tuned by the diffusion rate. For a small range of diffusion rates networks evolve the largest variety of topologies: modular, centralized or intermediate. Weighted networks in the transition range show topological but not weighted rich-club structure matching empirical data in the human brain. The simulation results allow us to propose adaptive rewiring based on diffusion as a parsimonious model for activity-dependent reshaping of the brain’s connections.

**Author Summary:** The brain is adapting continuously to a changing environment by strengthening or adding new connections and weakening or pruning existing ones. This forms the basis of flexible and adaptable behaviors. On the other hand, uncontrolled changes to the wiring can compromise the stability of the brain as an adaptive system. We used an abstract model to investigate how this basic problem could be addressed from a graph-theoretical perspective. The model adaptively rewires an initially randomly connected network into a more structured one with properties akin to the human brain, such as small worldness and rich club structure. The adaptive changes made to the network follow the heat diffusion, an abstract representation of brain functional connectivity. Moreover, depending on a parameter of the model, the heat diffusion rate, either modular or centralized connectivity patterns emerge, both found across different regions of the brain. For a narrow range of intermediate heat diffusion rates, networks develop a full range from modular to centralized connectivity patterns. Once settled into a connectivity pattern networks with normally distributed weights lock into that state, whereas networks with lognormally distributed weights show greater flexibility to adjust, while maintaining their small-world and rich club properties. Networks with lognormally distributed weights, therefore, show the combination of stability and flexibility needed to address the fundamental requirements of adaptive networks.

## Introduction

From neuronal synapses to white matter tracts (1), brain anatomical networks are characterized by structural properties like small-worldness (2), modularity (3, 4) and rich club organization (5). The pervasiveness of these properties raises the question whether they result from a common principle (4, 6).

We proposed that these properties are a product of adaptive rewiring (7–11, for a review 12). Adaptive rewiring captures a crucial property of how the brain anatomical network is shaped over time. The mechanisms that shape the network show great variety, as they encompass brain growth, development *and* learning (13 for a review). Yet they are alike in their common dependency on the network’s *functional connectivity*, i.e. the statistical dependencies between the nodes’ activities (10, 14). Adaptive rewiring formalizes this dependency in terms of graph theory, as it promotes adding shortcut links to regions of the network with intense functional connectivity, while pruning underused ones. These dynamical rewirings could be regarded as adaptive network optimization to function (15).

Adaptive rewiring, it turns out, robustly drives random networks to complex architectures matching the signature characteristics of brain anatomy; small world structure with a mixture of modular and centralized topologies. Adaptive rewiring models reach such “brain-like” structure, regardless of the specific choices made in representing the functional connectivity: rewiring works equally for oscillatory (7) or spiking activity (16), or for more abstract representations in terms of graph diffusion (9). Hence, adaptive rewiring may capture the common driving principle leading to such structure in the brain.

The robustness of ‘brain-like’ network evolution in adaptive rewiring models brings about a version of the well-known stability-plasticity dilemma (17). Adaptive rewiring involves a balancing act between two seemingly contrasting attributes: stability and flexibility. Whereas the robustness of adaptive rewiring warrants the stability of brain-like architectures, brains need also to be flexible enough to adapt to volatile circumstances. What are the conditions that enable flexibility in models that robustly develop ‘brain-like’ structure? Here we address this issue, which will involve a development of Jarman’s and colleagues (9) most abstract, and hence most general, adaptive rewiring model.

## Model

Recently, computational studies have shown that a simple, linear model can reliably predict functional from anatomical connectivity (18). In these studies, random walks represent the traffic through a fixed anatomical network. This allows the amount of traffic to be stochastically approximated in terms of graph diffusion. Jarman and colleagues (9) showed that adaptive rewiring shapes ‘brain-like’ networks, even when functional connectivity is essentially a matter of random walks on a graph.

An important shortcoming of Jarman’s and colleagues (9) adaptive rewiring model is that it uses binary networks to represents brain anatomy. Notwithstanding the popularity of binary graphs in brain network analysis, networks with differentially weighted connections offer a more appropriate representation of the sophisticated data on brain connectivity from contemporary tract-tracing and other imaging studies (2). Adaptive rewiring of differentially weighted networks does not automatically guarantee robust convergence to small world topologies; doing so may be ineffective since low-weight connections may not impact the network flow, whereas highly weighted ones may resist pruning (19).

Here, we first probe whether graph diffusion can drive the evolution of brain-like connectivity structures in weighted networks. Graph diffusion, we show, allows adaptive rewiring to establish ‘brain-like’ network structure, at least for two different weight distributions: normal and lognormal. The distribution of presynaptic weights on single neurons has traditionally been modeled as normal, but recent studies favor lognormal distributions, exhibiting a long tail of strong connections (20, 21). Our results show that both normal and lognormal weight distributions provide robust evolution to small world structure. Weighted networks also reproduce a key feature observed by Jarman and colleagues (9), involving the crucial character of *diffusion rate,* a parameter that represents in the model a global homeostatic or normalizing regulation mechanism (for a review 22). Diffusion rate is an important control parameter for the model: low diffusion rates give rise to modular structure, and higher ones to centralized ones.

In the intermediate range of diffusion rates there is a transition zone, in which networks show the greatest variability and complexity. After rewiring, both normal and lognormal networks at the transition zone can have modular or centralized structures. For normally distributed weights the network is equally likely to be modular, centralized or anything in between; for lognormal weights, the network are mostly in an intermediate state with both centralized and modular characteristics. In addition, models with both lognormally and normally distributed weights in the transition zone develop realistic rich-club structures, akin to backbone subnetwork structures in the human brain (23).

We find, moreover, that the transition zone endows adaptive rewiring with the requisite flexibility; i.e., for networks with lognormal but not with normal weight distributions. When the latter settle into either the modular or centralized state, they reach a ‘point of no return’. Networks with lognormal weight distributions, by contrast, can cross over from modular to centralized under continued rewiring. Outside of the transition zone, normally weighted networks continue to be in a ‘point of no return’ mode, whereas lognormally weighted networks are steered to a modular or centralized state (depending on whether the diffusion rate is below or above the transition zone), irrespective of their initial connectivity state. This combination of plasticity and stability, along with tuning by a single parameter, offers a solution to the stability-plasticity dilemma in the context of adaptive rewiring.

## Results

### Small-worldness

Small-worldness (S) is observed for all rewired networks, albeit with non-identical profiles across the system parameters τ and p_random_ (Fig 1A). S develops faster as a function of τ for binary networks than for normal and lognormal ones, the latter being the slowest to increase (Fig 1B). For τ = τ_plateau_ (different for each of the three weighting regimes), S reaches a maximum value, S_max_, henceforth sustaining it. S_max_ decreases significantly for p_rand_ >0.4. (Fig 1B).

**Fig 1.**
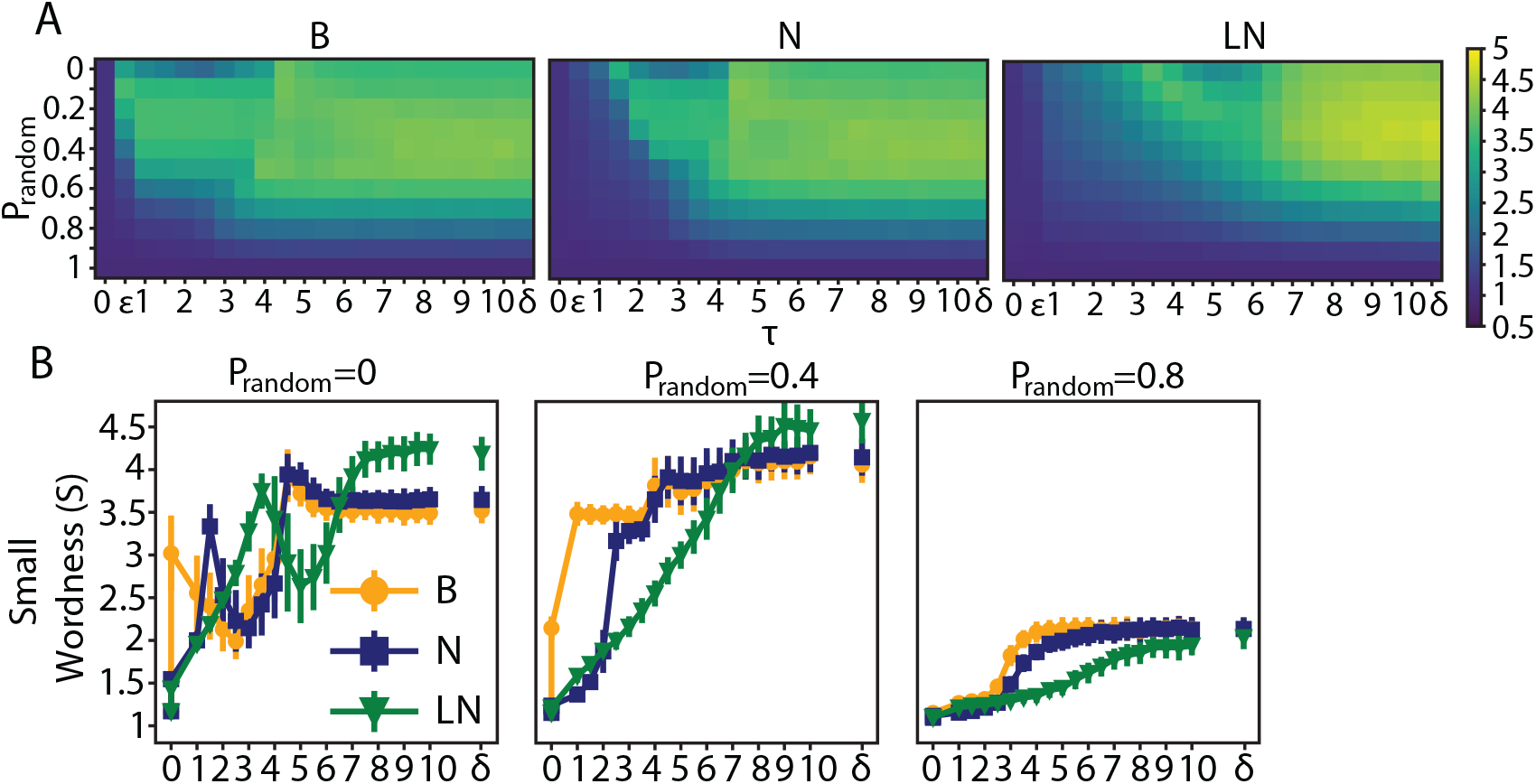
Networks of binary, normal and lognormal weight distributions develop into small worlds. A. From left to right, small world values (S) for networks with binary, normal and lognormal weight distributions for different heat diffusion rates (τ) and random rewiring probabilities (p_random_). ε= 10^-15^, δ = 10^15^. B. S as a function of τ for different p_random._ Horizontal bars indicate standard deviation.

An increase in a network’s average clustering coefficient (C) or a decrease in its average path length (L) induces by definition an increase in its small worldness. We probed the dynamics of C and L that drove up the small worldness of the networks. We found that along τ, S shows a strong positive correlation with C (ρ_binary_ = 0.91, ρ_normal_ = 0.96, ρ_lognormal_ = 0.97) and a moderate one with L (ρ_binary_ = 0.42, ρ_normal_ = 0.52, ρ_lognormal_ = 0.31). In the Watts and Strogatz model (24), the authors start with a structured network and parametrically swap ordered connections with random ones. This induces a significant decrease in L but a more moderate one in C, resulting in an increase in the small worldness of the rewired network. Our model is in a way the reverse process, in which rewiring induces a random network configuration to morph into a more structured one, and thus becoming small world due to an increase in C (See S1 Appendix for a more detailed description).

### Modularity

Small worldness characterizes networks with widely diverging topological qualities. For example, both of the adjacency matrices on the left and the right side of Fig 2 have the same S value. However, the ones on the left consist of clusters with dense intragroup and sparse intergroup connections. This characterizes modular connectivity patterns. On the other hand, the networks on the right comprise of a small number of nodes that are heavily connected, acting as hubs with the rest having very few connections. This is a feature of centralized topologies. Newman’s (25) modularity index (Q) differentiates between the networks on the left and right side of Fig 2 (Q values for the left vs. right network: Figue 2A, Q_binary_: 0.72 vs. 0.19; Fig 2B, Q_normal_: 0.70 vs. 0.22; Fig 2C, Q_lognormal_: 0.62 vs. 0.14).

**Fig 2.**
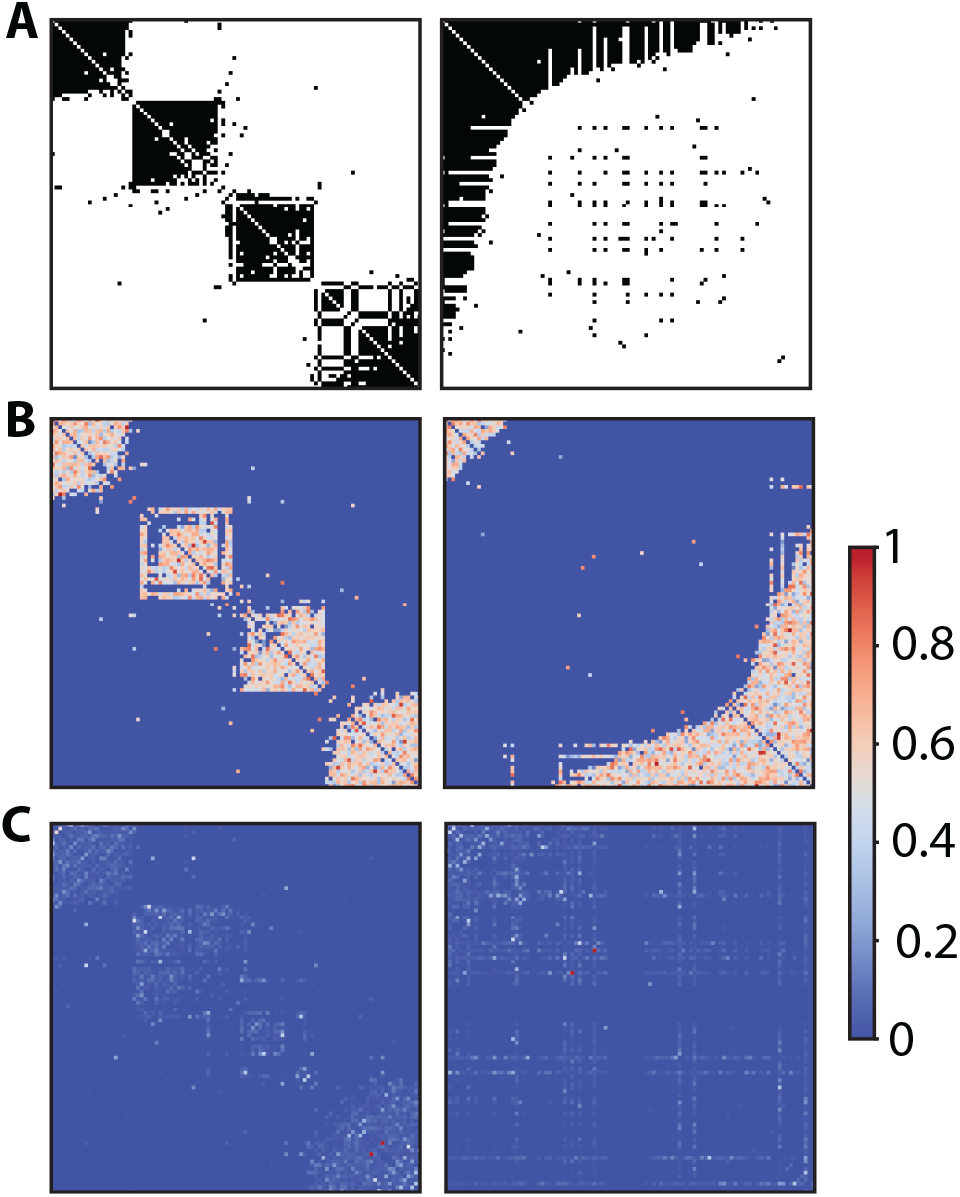
Networks can have the same small world value but diverging connectivity pattern. Adjacency matrices of small world networks with binary (A), normal (B), and lognormal (C) weight distributions. The adjacency matrices on the left are examples of modular networks where nodes within clusters are densely interconnected but sparsely interconnected between clusters. The ones on the right are examples of centralized networks where there is a degree imbalance between nodes, with the vast majority of the nodes having very few connections and the rest being heavily connected, acting as hubs. The networks on the left and right have the same small world value (A. S = 3.6; B. S=3.4; C. S = 2.3) despite their different topological characteristics. In all cases p_random_ is 0.2. For A. The values of τ for the left and right networks are equal to 2 and 5 respectively, for B. 3 and 5, and for C. 4.5, and 7.

Binary, normally and lognormally weighted networks develop modularity as a function of τ and p_random_ in a similar fashion (Fig 3A). Generally, Q initially increases as a function of τ, reaching a plateau, but eventually drops off to a value close to that of a random network (τ = 0) (Fig 3B). Binary networks reach their maximum Q faster (τ=10^-15^) compared to normal (τ=2), and lognormal networks (τ = 4) (Fig 3B). Furthermore, the binary networks sustain the maximum Q value for a greater range of τ values compared to the two weighted regimes, with the lognormal networks having the smallest range. For all networks, random rewiring diminishes the maximum Q value, with the binary networks being the most resistant and the lognormal ones the most exposed to this effect (Fig 3B).

**Fig 3.**
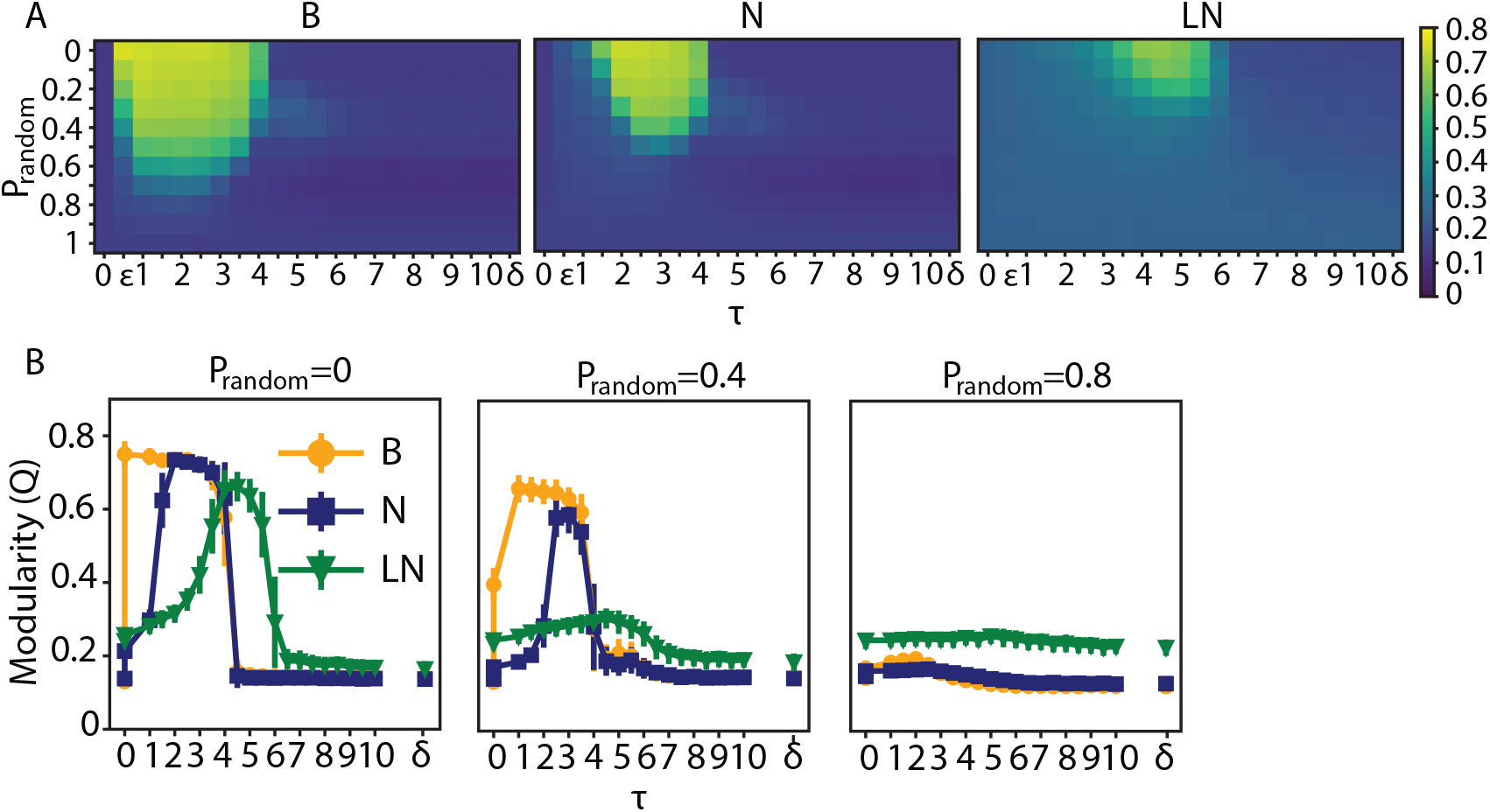
Modularity profiles between networks with binary, normal and lognormal weight distributions have similar shape. A. From left to right, Modularity index Q for networks with binary, normal and lognormal weight distributions for different values of the system parameters diffusion rate τ and proportion random rewirings p_random_. B. Q as a function of τ for different p_random_.

### Degree and strength distributions of modular and centralized networks

For a fixed number of total connections, centralized networks can be distinguished from modular ones according to the number of nodes with degrees deviating significantly from the mean. Centralized networks have a small number of high-degree nodes acting as hubs and, correspondingly, a large number of nodes with low degree. Both of these groups constitute outliers in the connectivity distribution. Hence, the proportion of outliers characterizes the centralization in the network. Degrees in the vicinity of the mean were established as <k> ± 3σ_κ_, where <k> is the mean degree of the network and σ_κ_ its dispersion. We used the Poisson distribution with mean <k> to calculate the value of the dispersion parameter (σ_κ_ = <k>^1/2^). A Poisson distribution is a suitable baseline distribution since random networks by virtue of their construction have most node degrees close to the mean and are well approximated by it. We probed how the proportion of outliers changes with τ, and which range of τ gives rise to centralized networks. We found that the proportion of outliers in networks as a function of τ follows a sigmoid function, with binary and normal networks having almost identical values but lognormal ones having a more gradual transition (Fig 4A).

**Fig 4.**
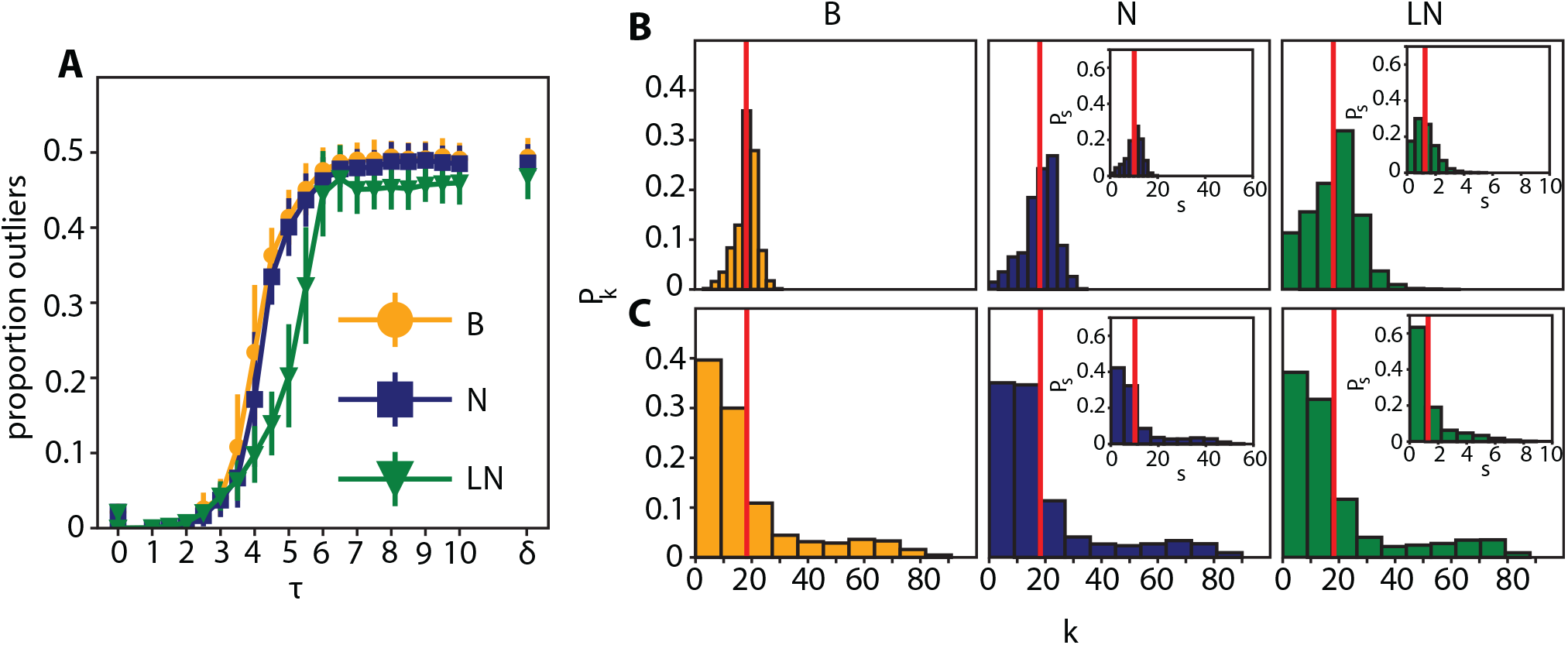
Degree distribution for modular networks is close to the mean value; for centralized networks it is heavy tailed. A. Proportion of nodes with outlier degrees as a function of τ for binary, normal and lognormal networks. B. From left to right, the distribution of degrees for binary, normal and lognormal networks (τ_binary_ = 2, τ_normal_ = 3, τ_lognormal_ = 4.5) which are modular. Inset plots show the strength distributions. C. Same as in B, but for centralized networks (τ_binary_ = 5, τ_normal_ = 5, τ_lognormal_ = 7). In all cases p_random_ = 0.2

Both the modularity and outlier analyses are in accord with each other: the range of τ values for which the networks have a small number of outliers includes the τ values for which the Q value is at its highest (it also includes τ values close to zero for which the Q value is small where the network is in a state starting to deviate from randomness but not quite reaching a definite structure). For τ values greater than this modular range, networks qualify as centralized according to our outliers criterion. Modular and centralized networks have different degree distributions, the former’s being concentrated around the mean (Fig 4B) and the latter’s having a large spread with a heavy tail (Fig 4C). The τ and p_random_ parameters giving rise to the distributions in Fig 4B are the same as in the example networks on the left side of Fig 2, for Fig 4C they are on the right side of Fig 2. Furthermore, weighted networks show the same degree and strength distributions (Fig 4B and 4C, inset plots).

We probed the heavy tail degree and strength distributions of representative centralized networks (τ_binary_ = 5, τ_normal_ = 5, τ_lognormal_ = 7, in all cases p_random_ = 0.2) The power-law and lognormal distributions were a better fit than the exponential one (Fig S2). Power law distribution functions are of the form P(k) ∼ k^-α^. For the degree distributions the exponent α that fitted best the data was 1.7, for the strength distributions it varied between 2 and 2.6 for the different types of networks (See S2 Appendix for a more detailed description of the analysis). We used publicly available code for this analysis (26).

### Network structure at the transition range

Our previous analyses showed that, for a given random rewiring probability, and depending on the value of the control parameter τ, the network could be rewired to be either in a modular or centralized state. At the boundary between those two states the structure of the network is ambiguous. Moving the control parameter τ across the boundary causes a phase transition from modular to centralized connectivity. Our aim in this section is to probe the properties of the network at this transition.

We estimated the τ value in the middle of the phase transition from modular to centralized networks (τ_transition_) by locating the point with the largest derivative value on each of the sigmoid curves in Fig 4A (inflection point). The inflection point is similar for both binary (τ_transition_ = 4.1) and normal networks (τ_transition_ = 4.15) but shifts to the right for lognormal ones (τ_transition_ = 5.5). In all cases, τ_transition_ gives rise to networks with the highest variability of outlier values compared to all the other τ values. In a similar analysis on the modularity data we found the same τ_transition_ values (finding the highest absolute derivative value for the descending part of the modularity curves and the points with the highest variance).

Regarding the distribution of modularity values at τ_transition_, one hypothesis is that the distribution is bimodal with one peak centered at the modular region and the other at the distributed one. A contrasting hypothesis is that the distribution is unimodal with the peak in-between the modular and centralized regions. For both binary and normal networks, we found a compromise between those two hypotheses: the modularity distribution at τ_transition_ is uniform, with the modularity values on the left and right boundaries giving rise to modular and centralized networks respectively (Fig 5A and B, center plots). Furthermore, depending on the sign of small perturbations (δτ = 0.1) to τ_transition_, the resulting modularity distributions show a distinct peak either at the modular region (for τ = τ_transition_ -δτ) or the centralized one (for τ = τ_transition_ +δτ) (Fig 5A and B, the leftmost and rightmost plots). On the other hand, the modularity distribution for lognormal networks is more in line with the second hypothesis: it is unimodal, and its peak is in-between the modular and centralized regions. Unlike the other two networks, its peak is not shifted for the small perturbations of its τ value (Fig 5C).

**Fig 5.**
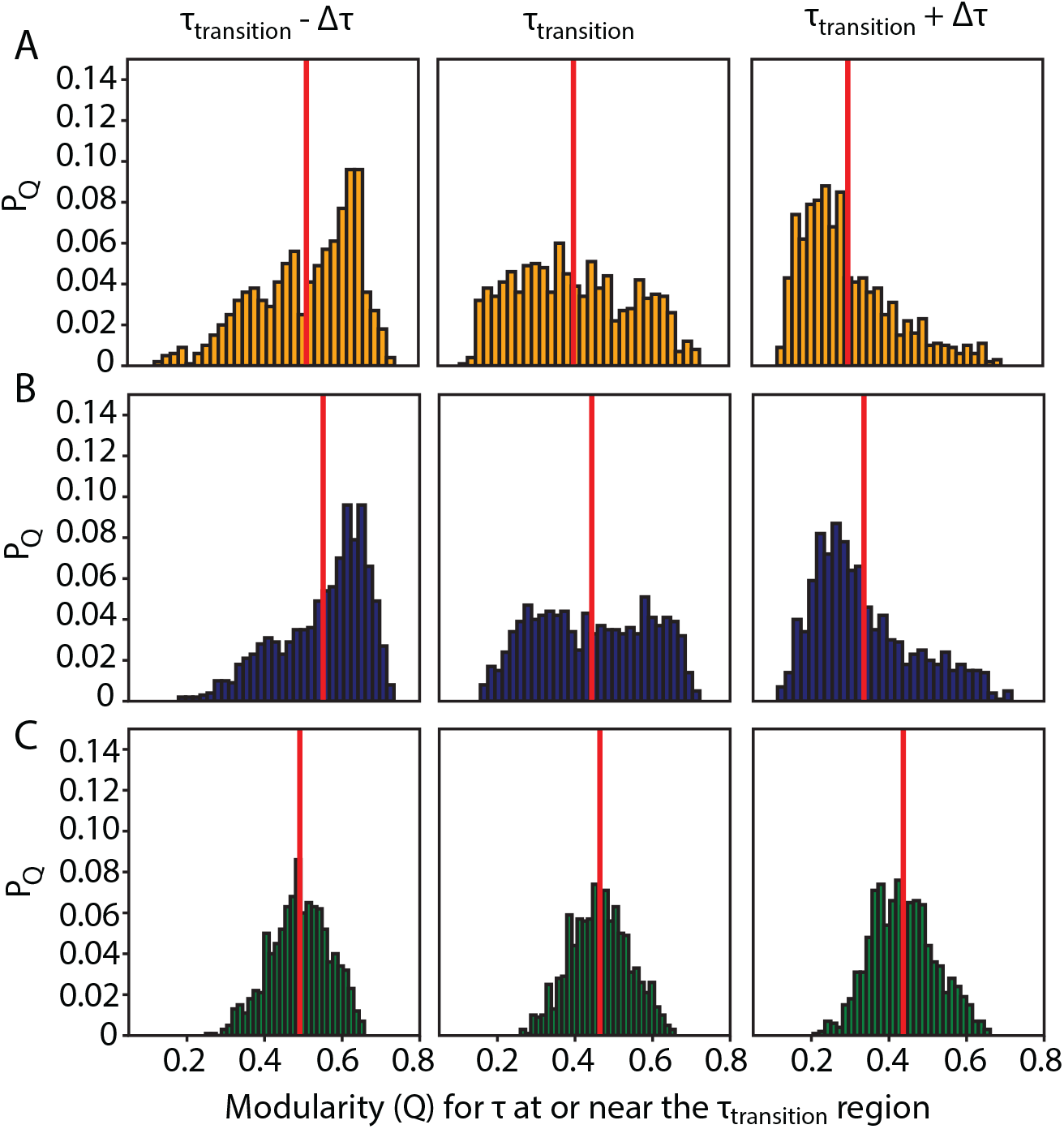
Binary and normal networks have a uniform modularity distribution and are more prone to connectivity transitions after slight perturbations to τ_transition_ compared to lognormal ones which have a modularity distribution with a distinct peak. A. Binary network. From left to right, modularity distributions for τ = (τ_transition_ -δτ, τ_transition_, τ_transition_ + δτ) = (4.0, 4.1, 4.2). For each τ we obtained 1000 modularity values each corresponding to a different instantiation of the rewiring algorithm B. Same as A for the normal network τ = (4.05, 4.15, 4.25). C. Same as A for the lognormal network τ = (5.4, 5.5, 5.6).

Starting from a random configuration, the rewiring process at τ_transition_ can result in vastly different connectivity patterns ranging from highly modular to highly centralized networks (Fig 5). On the other hand, at the fringes of its variability range, the initial random network is biased towards a more modular or centralized connectivity pattern. We asked whether small biases in the initial random network are amplified during rewiring, essentially predicting the connectivity pattern of the final network. We tested this hypothesis by comparing the modularity indices of 1000 networks before and after rewiring. We found a very weak correlation between the modulation values of the random and rewired networks (ρ_binary_ = 0.10, ρ_normal_ = 0.14, ρ_lognormal_ = 0.08; Fig 6A). This indicates that the connectivity pattern of the rewired network at τ_transition_ is nearly independent of the connectivity bias of the initial random network.

**Fig 6.**
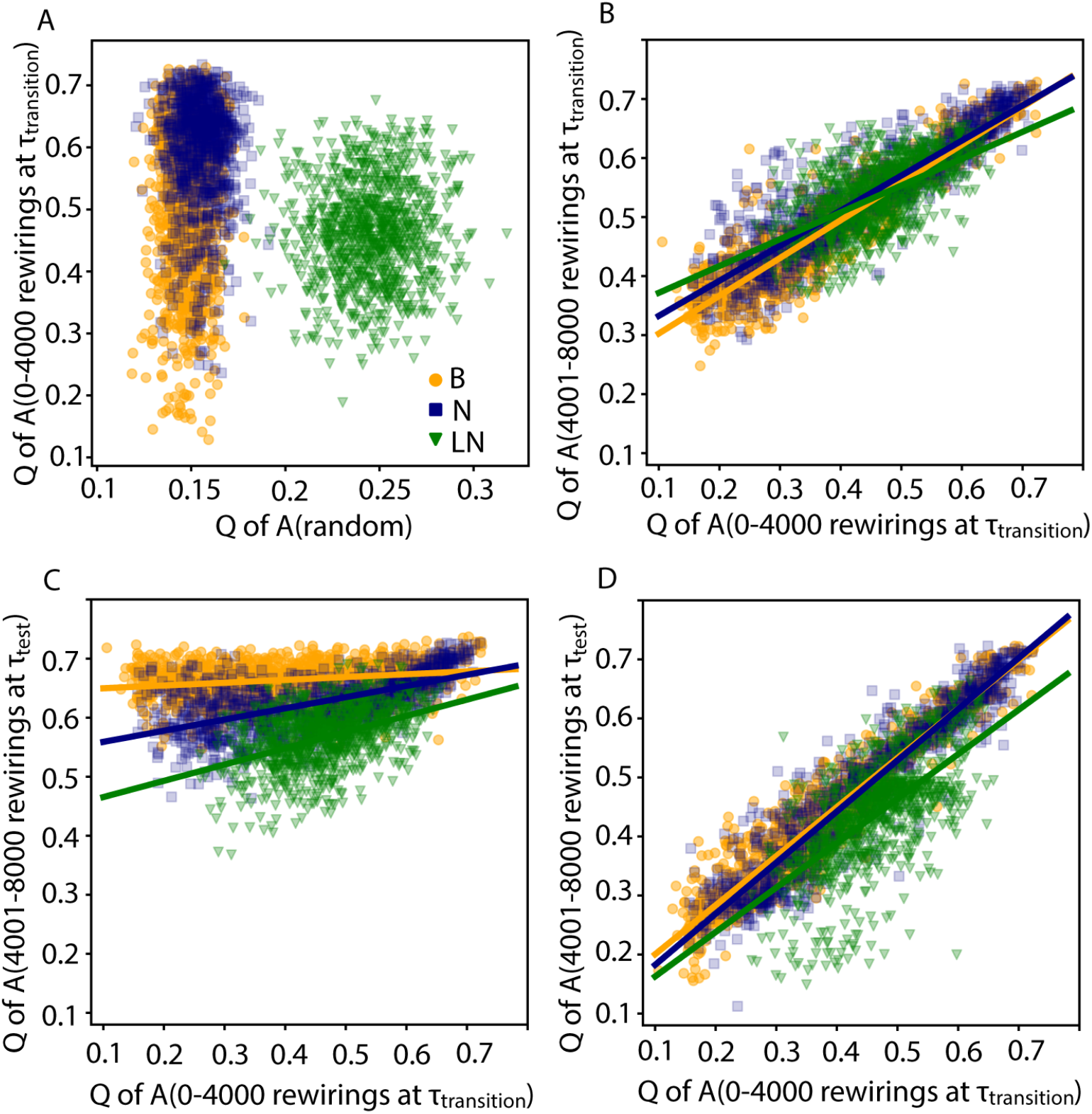
Flexibility and locking-in of networks vary depending on the weight distribution of their edges. A. Scatter plot showing very weak correlations between the modularity values Q of networks after 4000 rewirings at τ_transition_ plotted against those of their initial random configuration. B Scatter plot showing a strong positive correlation of the modularity values between same networks after 4000 and 8000 rewirings at τ_transition_. C. Same as (B) but now between 4000 and 8000 rewirings the τ values equal ones producing modular topologies from random initial configurations (τ_binary_ =2, τ_normal_ = 3, τ_lognormal_ = 4.5) D. Same as (C), but with τ values equal to ones producing centralized topologies from random initial configurations (τ_binary_ =5, τ_normal_ = 5, τ_lognormal_ = 7). The lines in all cases indicate the best linear fits to the data.

To observe whether established network connectivity patterns lock the rewiring at τ_transition_ in its configuration, we compared the modularity values of the final networks from the previous analysis (4000 rewirings) with the modularity values after 4000 additional rewirings (i.e. after 8000 rewirings in total). For both binary and normal networks we observed very strong correlations (ρ_binary_ = 0.94, ρ_normal_ = 0.89; Fig 6B) between the modularity values after 4000 and 8000 rewirings, indicating ‘a point of no return’: that is, once the network settles to a connectivity state, further rewiring no longer steers it away from that state. The correlation for lognormal networks is significantly less than the other two types (ρ_lognormal_ = 0.59; Fig 6B). This difference is also observed in the best (in the least squares sense) linear fits of the scatter plots in Fig 6B. The slope of the line for the binary (0.64) and normal networks is greater than the one for the lognormal (0.45) ones (Fig 6B). These results indicate that lognormal networks are more prone to change their connectivity state, most likely due to the small fraction of connections having significantly larger weights compared to the rest. Thus, in the sense that they do not lock to a specific state at τ_transition_, lognormally weighted networks show considerable flexibility.

Our next analyses probed the flexibility of network developed at τ_transition_, when τ is subsequently set to a different value. An initially random network was rewired 4000 times at τ_transition_; next another 4000 rewirings were subsequently performed at τ_test_. As long as τ_test_ deviates from τ_transition_ only slightly (τ_test_ = τ_transition_ ± 0.1), the network’s final connectivity pattern remains close to the one obtained after the first 4000 rewirings (more so for normal and binary networks than for lognormal ones; Fig S3). Fig 6C shows that with τ_test_ << τ_transition_, all networks are driven to modular connectivity patterns irrespective of their initial topologies. On the other hand, for τ_test_ >> τ_transition_, network evolution is strongly affected by the state originally reached at τ_transition_, at least for the binary and normal networks (Fig 6D). For lognormal networks, however, the correlation coefficient is low, indicating that the explanatory strength of the linear fit is weak.

For small τ_test_ values (τ_test_ < τ_transition_) all types of networks show similar behavior; the greater the difference between τ_test_ and τ_transition_ the less dependent the rewired network’s topology under τ_test_ from its initial topology (Fig 7A, correlation values for τ_test_ - τ_transition_ < 0). The same is true for τ_test_ - τ_transition_ > 0 but only for lognormal networks: as τ_test_ grows further from τ_transition_ the correlation in their topologies (modularity) decreases. Therefore, for lognormal weights, as the diffusion rate strays away from τ_transition_ in either direction, a network’s topology after rewiring becomes less dependent on its initial connectivity state. However, this is not true for binary and normal networks for larger τ_test_ values (τ_test_ - τ_transition_ > 0): the correlation value does not diminish as the difference between τ_test_ and τ_transition_ increases (Fig 7A), indicating that for large diffusion values the topology of rewired normal and binary networks depends on their initial configuration (Q_transition_).

**Fig 7.**
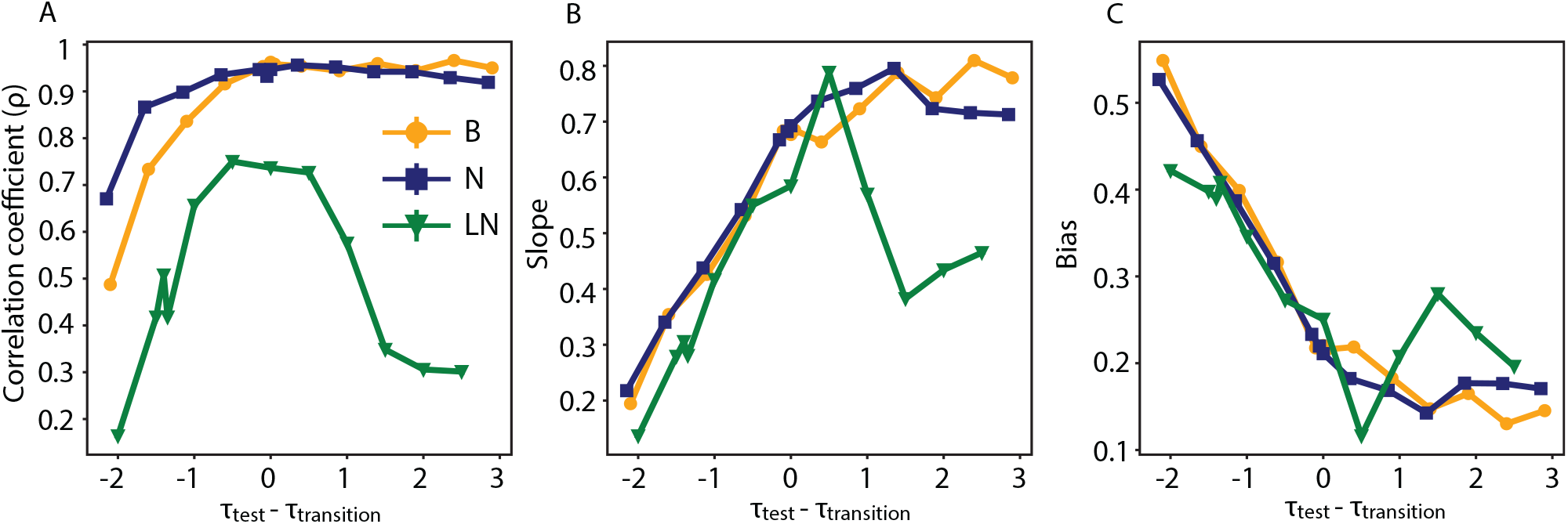
Lognormal networks show different flexibility patterns compared to normal and binary networks, especially for τ_test_ > τ_transition_. A. Correlation coefficient of the modularity index between the same network that was initially rewired at τ_transition_ (4000 rewirings) and then re-rewired at different τ_test_ values (an additional 4000 times). Each correlation point is for 100 iterations. C. and D. The slope (C) and the bias (D) of the best-fitted line that explains the data obtained from A.

We found the best linear fits for the modularity data produced for each pair of τ_test_ and τ_transition_ parameters (Fig 7B and 7C). More specifically, the linear fit takes the form Q_test_ = αQ_transition_ +β, where α and β are two scalar terms, the slope and the bias that we adjust to find the best fit of the data (Q_test_, Q_transition_) in the least squares error sense. The linear fits further cemented our findings from the correlation analysis. For small τ_test_ values (τ_test_ < τ_transition_) the best fit parameters show the same trend for all types of networks: the further away τ_test_ is from τ_transition_ the smaller the slope and the greater the bias. This indicates that for τ_test_ << τ_transition_ the rewired network will settle into a topological state that depends more on the bias term found from the fit (in the modular regime) than from its initial topological configuration (Fig 7B and 7C). For large τ_test_ values (τ_test_ > τ_transition_), the binary and normal networks sustain both the slope and bias values that the network has when τ_test_ = τ_transition_, but the lognormal networks do not, showing a tendency to decrease in slope and increase in bias as τ_test_ increases, an approximate mirror image of their behavior for small τ_test_ values (Fig 7B and 7C). Thus for τ_test_ > τ_transition_, binary and normal networks lock in their initial connectivity pattern similarly to when τ_test_ = τ_transition_, whereas the lognormal network shows a more flexible behavior with a bias nevertheless to switch into a more centralized state.

### Assortativity and rich club structure

We probed the possibility that the rewiring process favors homophily by measuring the topological assortativity coefficient, r. Networks with positive r are assortative, meaning that nodes with similar degrees tend to connect. Networks with negative r are dissasortative; in this case nodes with dissimilar degrees are more prone to connect. We found that for modular networks, r shows weak or zero assortativity; centralized networks dip into the dissasortative realm (Fig S4)

We further measured the rich club coefficient, Φ(k), of the networks. Topological rich club refers to the tendency of typically high degree nodes to connect with each other; when its normalized counterpart, Φ_norm_(k), is above the baseline (greater than 1), then the subset of nodes with degree greater than k is more densely interconnected compared to a random control with the same degree distribution. The coefficient is not trivially connected to assortativity, since a dissasortative network could still be rich club and vice versa (27). We found that in the modular and transition range the weighted networks exhibited topological rich club behavior for a range of intermediate degrees (binary networks were not rich club for τ_transition_), whilst centralized networks were not rich club for any degree (Fig S5).

For normal and lognormal networks, we evaluated the normalized weighted rich club coefficient, Φ_w,norm_(k). Φ_w,norm_(k) above the baseline indicates that the edges of the rich club nodes have larger weights compared to a random control network (a network with the same topology but randomly reshuffled weights). We found that for the lognormal networks Φ_w,norm_(k) did not deviate from 1; for the normal networks, Φ_w,norm_(k) similarly hovered around 1 in the modular and centralized regimes. Hence for these states the rewiring process does not distribute the larger weights preferentially to the high degree nodes. In general the data show that there is a distinction between the topological and weighted measures; the former being above baseline for a range of degrees, but not the latter.

This distinction becomes more accentuated for normal networks in the transition zone, where Φ_w,norm_(k) tilts below baseline for nodes with larger degrees (k>20) (Fig 8). Taken together, these topological and weighted coefficient profiles bode well with anatomical data on the human brain. Specifically, van den Heuvel and colleagues (23) mapped human brain structural connectivity by estimating via diffusion tensor imaging (DTI) the number of connecting white matter streamlines between 1,170 subdivisions of the cortex. Based on topology the human network exhibits rich club behavior, but its weighted counterpart does not (from 28 Fig 3b and 3c), exhibiting the same qualitative properties as the normal –and the lognormal to a lesser extent-network at τ_critical_.

**Fig 8.**
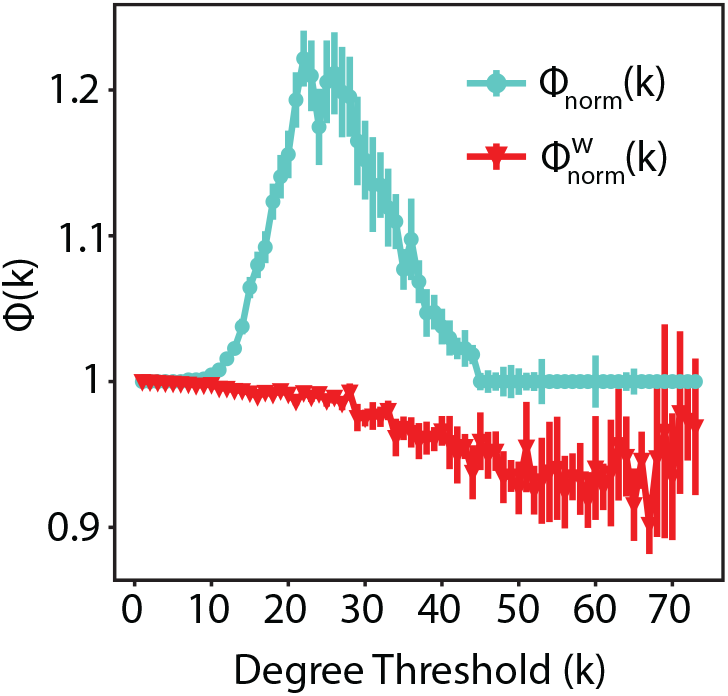
Topological and weighted rich club coefficients diverge similarly to physiological data. Topological and weighted rich club coefficient for the normal network at τ_transition_ = 4.15, p_random_ = 0.2

We performed a fine-grained analysis at τ_transition_ to probe the rich club characteristics of networks with modular, centralized and in-between connectivity patterns. From the analysis data in Fig 5, we used the adjacency matrices corresponding to the 50 smallest modularity values to obtain the rich club measures for the centralized connectivity patterns, the 50 adjacency matrices corresponding to the largest modularity values for the modular patterns, and the 50 matrices corresponding to the modularity values between the mean one for the in-between patterns. The weighted networks showed rich club characteristics in all three cases (Fig 9A). Thus the difference between the centralized networks at τ_transition_ and at τ > τ_transition_ is that in the former case the network is rich club (for 20<k<40), in the latter it is not. This effect is more accentuated for the lognormal network. Visual inspection of the adjacency matrices of the different connectivity patterns at τ_transition_ indicate a smooth transition from modularity to centrality (Fig 9B and 9C).

**Fig 9.**
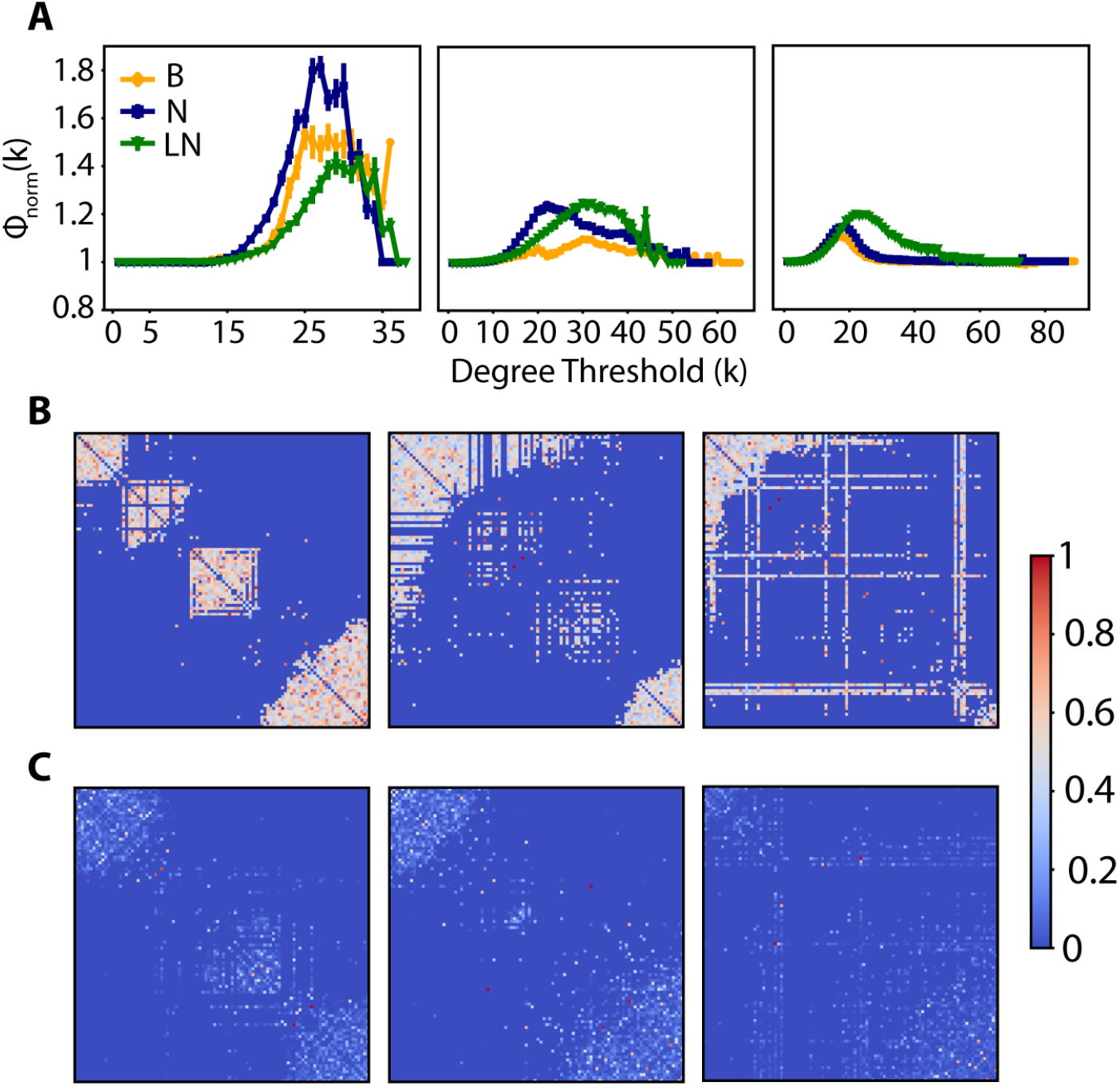
Weighted networks are rich club for all the topological subdivision at τ_transition_. A. Topological normalized rich club, Φ_norm_(k), for binary, normal and lognormal networks in the modular (left), centralized (right), and in between (center) parts of τ_transition_. B. Example adjacency matrices of normal networks in the modular (left), centralized (right), and in between (center) parts of τ_transition_. C. Same as in B but for lognormal networks.

## Discussion

### Graph diffusion and other rewiring models

We showed how in graph-theoretical terms, the stability-plasticity dilemma can be addressed in adaptive rewiring of weighted model networks. In these models, rewirings are made to bolster the functional connectivity. In the present model, functional connectivity was represented by heat diffusion. In previous adaptive rewiring models, functional connectivity was represented as synchronous activity (7,8,10,11,16,19,29,30) of linearly coupled nonlinear oscillators (31) or, somewhat more realistically, by spiking model neuron synchrony (16, 32).

While these representations more closely resemble artificial neural networks than the current graph diffusion model, their shortcomings are twofold. They rely on arbitrary functional connectivity dynamics, whereas a representation in terms of diffusion is empirically adequate (18). Moreover, rewiring in the older models involved a global search for the best (i.e. most synchronized) rewiring target. Even though this problem was remedied in Jarman and colleagues (29) by taking rewiring distance into account, this introduced arbitrary assumptions about the spatial embedding of these networks. Jarman and colleagues (9) offers a graph-based solution to this problem, based on the rate of heat diffusion relative to that of rewiring. The lower the diffusion rate, the narrower the network search for the best rewiring candidate; higher diffusion rates allow the search to be incrementally broader. Diffusion rate offers a natural control parameter, to determine whether the resulting network structure tends to be modular or centralized. The diffusion rate parameter may represent global homeostatic forms of plasticity regulating the network’s dynamics (for a review 22).

### Graph diffusion and small worldness

For a wide range of species from C. elegans to humans, neuronal connections have been shown to form small world networks, with small path length and high clustering coefficient (2,24,33). Adaptive rewiring based on diffusion when driven by the network’s own functional connectivity, leads to a similar small world structure. Networks are evolved to be small world for a wide range of diffusion rates and random rewiring probabilities (Fig 1). This is by and large because diffusion shapes the initial random network into a more structured connectivity pattern with high clustering coefficient while at the same time it maintains small path lengths (Fig S1)

Adaptive rewiring based on diffusion is an abstraction of the consequences of Hebbian learning, where connections with heavy traffic are strengthened whereas the ones with less activity are weakened or pruned. A similar process may lead to changes in the connectivity pattern of higher cognitive regions throughout childhood and adolescence. For example, using resting state functional connectivity MRI on children and adults Fair and colleagues showed that with age short-range functional connections between regions that are involved in top-down control regress whereas some long-range functional connections develop or increase in strength (34). This process leads the connectome of the regions involved mostly in higher cognitive functions to have a small world structure (35).

### Diffusion rate and network topology

Even though small-worldness is a key property of biological organs; many systems with similar small world value have diverging topologies perhaps because of differentially weighted trade-offs between multiple constraints (36). Diffusion-based models show this variety across the range of the diffusion rate parameter τ. Across a wide range of diffusion rates, adaptive rewiring leads to approximately the same small world values, but different topological structures (Fig 2). For smaller τ values the emerging network is modular (Fig 3), with dense connectivity within clusters but sparse between them. For larger τ values, we obtain centralized topologies where a subset of nodes are acting as hubs (Fig 4C). Both of these patterns are present in the brain. Clustered neuronal modules facilitate local computations, an example being cortical minicolumns, the highly structured functional patterns in the sensory cortex. Neurons within a minicolumn and typically across the layers are densely connected, but tangential ones, across the layers to other minicolumns are sparsely connected (37). On the other hand, centralized connectivity patterns resemble brain modules that receive and integrate a distributed set of neural inputs, for example the different association areas in the cortex (38, 39).

In the centralized regime, all types of networks show a similar degree distribution that is approximated by a power law distribution with an exponent of 1.7, for both binary and weighted networks (Fig S2). Diffusion tractography (40) and fMRI methods (41) have shown a degree distribution between cortical and subcortical regions that follow an exponentially truncated power law with exponents of 1.66 and 1.80 respectively. Brain functional connectivity inferred from fMRI in different tasks shows a scale free distribution with a power law exponent of 2 (42).

### Rich club structure in the transition zone

Brain networks at different levels and for different species have been shown to be topologically rich club (23,43,44), that is, regions or neurons acting as hubs are also preferentially connected among themselves. We found that the nodes of both the normal and lognormal networks show topological rich club behavior for small up to transition diffusion rates. Their corresponding weighted rich club index, which measures to what extent larger weights are attached to the connections within the rich club, was either at or below baseline (Fig 8). This interesting contrast between topological and weighted rich clubs is in line with physiological data on the human brain (23, 28).

Furthermore, the centralized structures in the transition zone showed rich club behavior (Fig 9), but the centralized networks for larger τ values did not (Fig S5). The combination of centrality and rich club behavior is a signature of connectivity patterns in the brain (5). Rich club connections in the brain could constitute its backbone, communicating information from diverse functional modules. Empirical evidence has indicated that rich club connections extend over long distances forming a high capacity structure that facilitates global communication (23).

### Flexibility and stability of topology in and out of the transition zone

In the transition zone between modular and centralized structures, we observe the highest variability in network topology. For example, in contrast to the τ values producing robustly either modular or centralized topologies-when starting from a randomly connected one-, at the center of the boundary between them, τ_transition_, a network’s connectivity pattern cannot be predefined before rewiring since it is equally likely that it will have either topology and anything in-between in a continuum (Fig 5). Whereas this transition zone is similar in range for networks with binary and normally distributed connection weights, it covers a considerably broader parameter range for networks with lognormally distributed weights (Fig 4A).

Very small connectivity biases in the initial random network do not influence the final topology of the rewired network (Fig 6A), however, larger biases lock the network at the connectivity state it already is (Fig 6B). This ‘point of no return’ is not observed for networks with lognormal weights. Moreover, lognormal networks show a more controllable behavior as the diffusion rate moves away from the transition range in either direction: for diffusion values smaller than the transition ones networks will most likely evolve to become modular and for greater values they develop into a centralized topology irrespective of their initial connectivity pattern (Fig 7).

Throughout these modifications, the networks retain their small-world characteristics. The combination of maintaining the small-world topology and its flexible malleability, controlled by a single parameter, is a desirable characteristic for learning mechanisms associated with plastic modifications in the brain (45–47). In lognormally distributed networks the emphasis on modularity versus centrality around τ_transition_ varies smoothly with τ. This smooth transition may also be considered advantageous for neuronal networks, since slight changes of a global regulating process will not have dramatic effects in their connectivity pattern. This combination of stability and flexibility adds to the advantages of lognormal networks mentioned elsewhere, such as sparsity in spontaneous firing rates (48) and efficient coding (49).

## Materials and Methods

### Graph preliminaries

A graph is defined as an ordered pair *G = (V, E, W)*, where *V* denotes the set of vertices (or nodes), *E* the edges (or connections) between them, and *W* the set of edge weights, 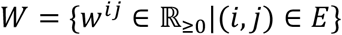, i.e. only nonnegative weights are used. The cardinalities, |*V*| = *n* and |*E*| = *m*, express the total number of nodes and connections respectively. The connectivity pattern of *G* can be conveniently represented by an n x n adjacency matrix, *A*, with its entries denoting the weights between edges (*A_ij_* = *w^ij^*). *w^ij^* = 0 signals that edges *i* and *j* are not adjacent. If *w^ij^* > 0 then vertices *i* and *j* are adjacent; the greater the value of the weight the stronger the connection between vertices. In the case of a binary network, the weights and the entries of the adjacency matrix can take only two values, 0 or 1; *A_ij_* = 1 indicates that vertices *i* and *j* are adjacent ((*i*, *j*) ∈ *E*), and *A_ij_* = 0 that they are not ((*i*, *j*) ∉ *E*). In this paper both binary and weighted graphs are undirected and simple (there are no self loops), meaning *A* is symmetric (*A_ij_* = *A_ij_*), and zero in its diagonal entries (*A_kk_* = 0) respectively. The strength of a vertex *j* is the sum of the weights from the edges incident to it. Practically the strength is found by summing the rows or the columns of the adjacency matrix: 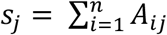. In the case of binary networks, the summation indicates the degree *d_j_* of node *j*.

### The graph Laplacian matrix

The graph Laplacian is defined as *L* = *D* − *A*, where D is a diagonal matrix having the strengths (or degrees for binary adjacency matrices) of the nodes in its diagonal entries (*D_ii_* = *s_i_*). It arises naturally in optimization problems such as graph partitioning (50) or nonlinear dimensionality reduction (51). It also emerges as the discrete analogue of the Laplace-Beltrami operator 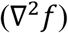 in the heat flow equation for graphs. The normalized graph Laplacian is defined as 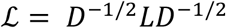, with 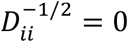 for *s_i_* = 0. Its entries take the values:

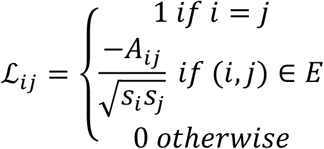

We use the normalized graph Laplacian, a more appropriate operator for graphs that are not regular, that is graphs with vertices that do not necessarily have the same strength (or degree). This is because the eigenvector of the normalized graph Laplacian corresponding to the zero eigenvalue captures the graph irregularity: 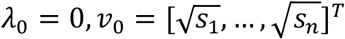 (52). Thus, throughout the paper, any mention of the Laplacian refers to the normalized version.

The Laplacian is a symmetric positive semidefinite matrix with real values: ℒ*_ij_* = ℒ*_ji_* and 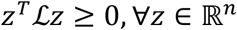; consequently, its eigenvalues are nonnegative and real and its eigenvectors form an orthonormal set. The spectral decomposition of the Laplacian matrix is ℒ = *V*Λ*V^T^*, where Λ = *diag* (*λ*_0_, *λ*_1_ … *λ*_n_) is a diagonal matrix, having in its diagonal entries the eigenvalues of ℒ (0 = *λ*_0_ ≤ *λ*_1_ ≤ ⋯ ≤ *λ*_n_ ≤ 2) and *V* = [*v*_0_*v*_1_ … *v*_n_] is an nXn matrix having as its columns the corresponding orthonormal eigenvectors (ℒ*v_i_* = *λ_i_v_i_*).

The eigenvalues and eigenvectors of the Laplacian yield valuable information about the graph they are derived from (53). The multiplicity of zero indicates the number of connected components in the graph; a single zero eigenvalue corresponds to a connected graph, two zero eigenvalues to a disconnected graph with two components, and so on. The second smallest eigenvalue, called the Fiedler value, and the largest one are related to several graph properties such as the graph’s connectivity, diameter and convergence to a stationary probability distribution for random walks (54–57). The values of the eigenvector corresponding the smallest nonzero eigenvalue, the Fiedler vector, can be used to sort vertices so that the ones close to each other belong to the same community, and accordingly partition a graph (50, 58). Nodes are sorted in the same order as the elements of the Fiedler vector in figures showing adjacency matrices.

### Heat kernel

The heat equation is a partial differential equation that describes how heat is spatially dissipated over time. It is defined as:

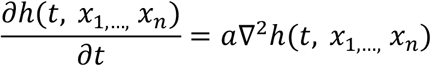

where *h* is a function of n spatial variables *x*_1_, …, *x*_n_ and time variable *t*, *a* is a positive constant and ∇^2^ is the Laplacian operator, or else the divergence of the gradient.

The heat equation based on the graph Laplacian matrix is similarly describing the variation in the flow of heat (or energy or information) within a graph over time:

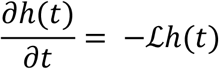

The heat kernel, *h*(*t*), and the graph Laplacian, ℒ, are both nXn matrices describing the flow of heat across edges and its rate of change respectively. *h*(*τ*)*_ij_* is the amount of heat transferred from vertex i to vertex j after time τ. The heat equation has an explicit solution:

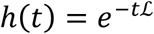

As it was shown in the previous section, ℒ can be decomposed to its eigenspectrum. The heat kernel can then be written as:

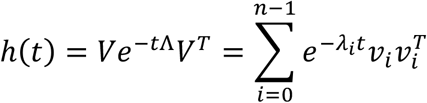

The dynamics for which diffusion spreads in the graph are based on the eigenvalue/eigenvector pairs of the Laplacian matrix. In the summation the contribution of each eigenmode is not the same, but depends on the magnitude of the eigenvalue. Typically, as t increases, only the eigenmodes with the few smallest eigenvalues play a significant role in the summation. For a connected graph, in the extreme case of infinite time (*t* = ∞), only the first eigenmode will contribute to the summation, since *λ*_0_ = 0.

### Adaptive rewiring algorithm

With the adaptive rewiring algorithm and its variations employed in this paper, we seek to probe the effects that a simple self-organizing rule will have on the properties of an initially randomly connected network. The core algorithm is essentially the same as the one in Jarman and colleagues (9), extended for application to both binary and weighted networks.

Starting with a random Erdös–Rényi type with |*V*| = *n* vertices and 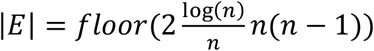 connections- the latter guaranteeing the random network is fully connected (59)- the rewiring algorithm proceeds as follows:

Step 1: Select, with uniform random probability, a vertex, *k* from the set of vertices in the graph that are of nonzero degree but also not connected to all other vertices 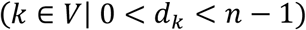.

Step 2: With probability p, go to step 2.1 (random rewiring), otherwise go to step 2.2 (heat diffusion rewiring). In both cases, select vertex *j*_1_ from the set of vertices that are not connected to *k*, 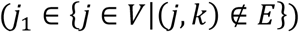, and vertex *j*_2_ from the set of vertices that are connected to *k*, 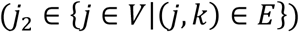, albeit in a differen. Delete the edge (*k*, *j*_1_), and add the edge (*k*, *j*_1_),. In the case of weighted networks, unless otherwise stated, use the weight of the previously connected edge (*k*, *j*_2_), for the edge (*k*, *j*_2_).

Step 2.1: For both *j*_1_ and *j*_1_ the selection is random and uniform among the elements of each set.

Step 2.2: Calculate the heat kernel, *h*(*τ*), of the adjacency matrix. *j*_1_ is selected such that from all the vertices not connected to k, it is the one with the highest heat transfer with k. *j*_2_ is selected such that from all the vertices connected to k, it is the one with the lowest heat transfer with k. In mathematical terms, we can express the statements above as follows:

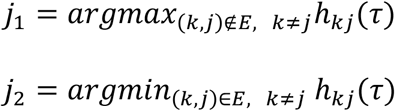

Step 3. Go back to Step 1 until k rewiring iterations are made.

### Graph measures

All the analyses were programmed in Python. The code package of the functions producing the rewired adjacency matrices, as well as the metrics described below characterizing them are publicly available (https://github.com/rentzi/netRewireAnalyze).

The *clustering coefficient*, C, measures the local cohesiveness of a network by quantifying the extent to which two nodes both adjacent to a common node are themselves connected to each other. The *weighted clustering coefficient*, C_w_, includes the weight of the edges forming these triplets. It is defined as:

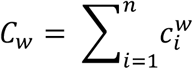

where 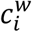 measures the clustering coefficient for node *i* and is expressed as:

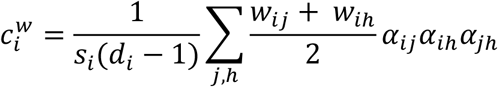

*s*_i_ is the strength and *d_i_* the degree of node *i*, *w_ij_* the weight of the connection between nodes *i* and *j*, and *α_ij_* a binary index that is 1 if *w_ij_* >0, and 0 otherwise (60). The formula above also works for a binary network.

A network is *small world* if it is connected in such a way so that the nodes connected to a common node are likely to be connected as well (high *C*), but at the same time any node can reach any other node in a few steps. The latter property refers to the average path length, *L*, which has important ramifications in a network’s behavior. For example, in a computational unit such as a visual module small *L* suggests that the results of local scene computations can be relayed fast to other components of the network for the computation of global effects (for example normalization). The metric that measures the small worldness of a network is defined as:

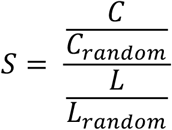

where *C* and *L* are the average clustering coefficient and path length of the network, and *C_random_*and *L_random_* are the ones for a random network with the same number of nodes and connections. A network with S>1 is considered small world.

To obtain a measure of network *modularity*, we used the spectral algorithm for community detection introduced by Newman (25), which identifies the communities within the network and also provides a modularity index (Q). The index Q gives a measure of the density of the interconnections within and between the found groups compared to a randomly connected graph with the same degree distribution: the greater Q is, the denser the connections within-and the sparser between-groups. For a division of a network into two clusters, we compute the eigenvector corresponding to the largest eigenvalue of the modularity matrix, *B*, and assign the nodes to one or the other group by the sign of the elements in the eigenvector. The entries of *B* are defined as:

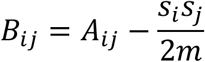

where *s_i_* is the strength of node *i*, and *m* is the total strength of the network. Q for this division into two groups is then:

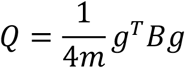

where *g_i_* is 1 is node *i* belongs to group 1, and −1 if it belongs to group 2. We iteratively continue to subdivide the groups and add the additional contribution ΔQ. We stop until any additional split will give us a negative ΔQ. For a complete description of the iterative algorithm see Newman (25).

A network is *assortative* if nodes with similar characteristics are preferentially connected. In our study the characteristic is degree. The assortativity coefficient (61) formally expresses this property as:

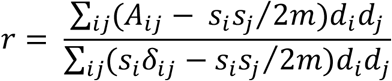

where A_ij_ is the weight of the connection between the *i* and *j* nodes, s_i_ is the strength of the *i*-th node, *m* is the total strength of the network and *d_i_* is the degree of node *i*. *r* can take values between 1 and −1, with 1 signifying a perfectly assortative network and −1 a perfectly disassortative one. The equation is generalizable for both binary (where *s_i_* is then equal to *d_i_*) and weighted network and is equivalent to the equations found in other studies for weighted networks (62, 63).

*Rich club* (27, 64) refers to high degree nodes preferentially connecting to each other. The topological rich club coefficient, Φ(*k*), disregards connection weights. It quantifies the tendency of the subgroup of nodes with degree greater than *k* to connect with each other. It is expressed as the ratio of the existing connections between the nodes with degree greater than k over the maximum possible connections between them:

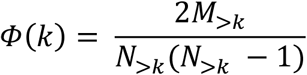

where N_>k_ is the number of nodes with degree greater than k, and M_>k_ is the number of connections between them. The rich club coefficient naturally increases with increasing k, even in randomly connected networks. We therefore normalize Φ(k), with the coefficient, Φ_random_(k), obtained from a random network with the same number of nodes, connections, and degree distribution as the original one. We construct the random network by performing 100*M_>k_ double edge swaps from the original network. A double edge swap involves randomly choosing two connections a-b and c-d, and replacing them with the connections a-c and b-d. If the later connections already exist, then the algorithm does not count that iteration and chooses again two random connections (43). The normalized rich club coefficient is expressed as:

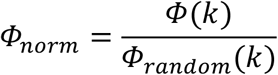

For the weighted rich club coefficient, Φ_w_(k), the node selection is based, similarly to Φ(*k*), on the degree k. Φ_w_(k) is expressed as:

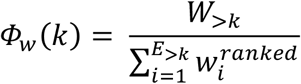

where W_>k_ denotes the sum of the connection weights between the nodes with degree greater than k, E_>k_ is the number of connections between them, and w^ranked^, the weights in the network in descending order. To normalize the weighted rich club coefficient, we use Φ_W,random_(k), which has the same topology as the original network but reshuffled edge weights (28, 65). The weighted normalized rich club is then:

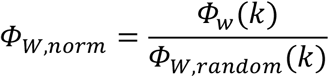

### Simulations and analyses

Simulations and analyses were performed on 100-node networks (average degree was 18.24). Most simulations vary two parameters from the rewiring algorithm: the random rewiring probability (p_random_) and the time variable in the heat equation (t = τ). For each combination of parameter values, we run the rewiring algorithm 100 times. Unless otherwise stated, the figures show mean values of these 100 runs, with the vertical lines at the measured points indicating standard deviation. Each run involves 4000 rewirings, since by then networks typically have diverged from a random connectivity pattern.

## Acknowledgments

We thank Steeve Laquitaine for inspiring discussions.

## S1 Appendix

We performed pairwise correlations (Pearson coefficient; ρ) along the τ dimension, separately for each of the 100 iterations and p_random_ bins (pairs of p_random_, 0.1 apart) to quantify the dependencies between path length (L), average clustering coefficient (C) and small-world (S) metrics. Qualitatively, C shows a similar pattern to S: steepest increase with respect to τ for binary networks, and increasingly moderate for the normal and lognormal distributions (Fig S1A). C is perfectly correlated with S (average across iterations and probability bins was ρ_binary_ = 0.91, ρ_normal_ = 0.96, ρ_lognormal_ = 0.97).

By contrast, L is not a monotonically increasing function of τ but reaches a peak and subsequently reverts back to values close to those of random connectivity networks; the peak is reduced to an almost flat line for *p_rand_* >0.4 (Fig S1B). Overall, across τ, L was positively correlated with C, for all networks (ρ_binary_ = 0.42, ρ_normal_ = 0.52, ρ_lognormal_ = 0.31). The correlation results along with the S values indicate that the diffusion rewiring increases small worldness by increasing clustering at a greater rate compared to path length, something that is more accentuated for larger random rewiring probabilities (compare Figue S1A with S1B).

**Fig S1.**
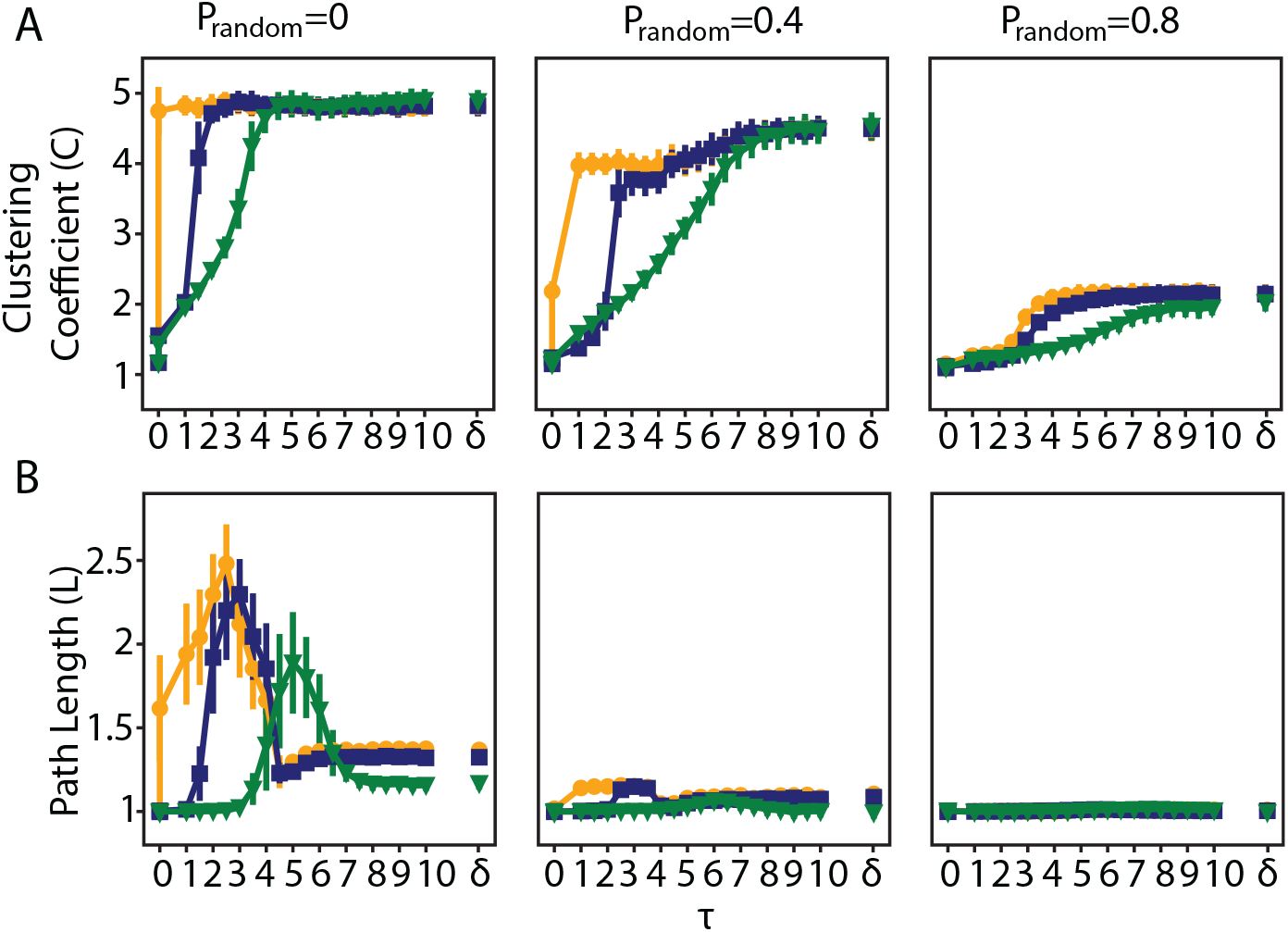
An increase in C as a function of τ is accompanied by a more moderate increase in L. A. C as a function of τ for different p_random._ (0, 0.4, 0.8). B. Same as A for the L metric. Horizontal lines indicate standard deviation.

## S2 Appendix

To probe the heavy-tailed degree distributions of the centralized networks, we used p_random_ = 0.2, and τ_binary_ = 5, τ_normal_ = 5, τ_lognormal_ = 7 as representative τ values for centralized networks. Using log-likelihood ratios, we determined whether they best fit a power-law, lognormal or exponential distribution. We first compared the lognormal with the power-law fit, and then the better fit of the two with the exponential distribution. We repeated this process for 100 different realizations of the network. In over 90% of the cases either power-law or lognormal were a better fit compared to exponential distributions, with no significant difference between lognormal and power-law. The fits were adjusted so that they optimally accommodate the tail of the distribution, i.e. if they did not improve the fit, bins with the smaller degrees were not included. For the optimal range of degrees at the tail end of the distributions, the lognormal and power-law fits were almost identical (Fig S2, fits on the left side of the plots). We performed the same analysis for networks of 1000 nodes (this time for one iteration due to the computational load) to probe whether the fit changes when the maximum allowable connections of a node increases by an order of magnitude with similar conclusions (but worse fits). Power law distribution functions are of the form P(k) ∼ k^-α^. For all networks the exponent α was the same up to a decimal (α = 1.7). The strength distributions of the normal and lognormal networks again showed a similar behavior to their degree counterparts with the only difference being that the distribution decayed faster (exponent α varied between 2 and 2.6).

**Fig S2.**
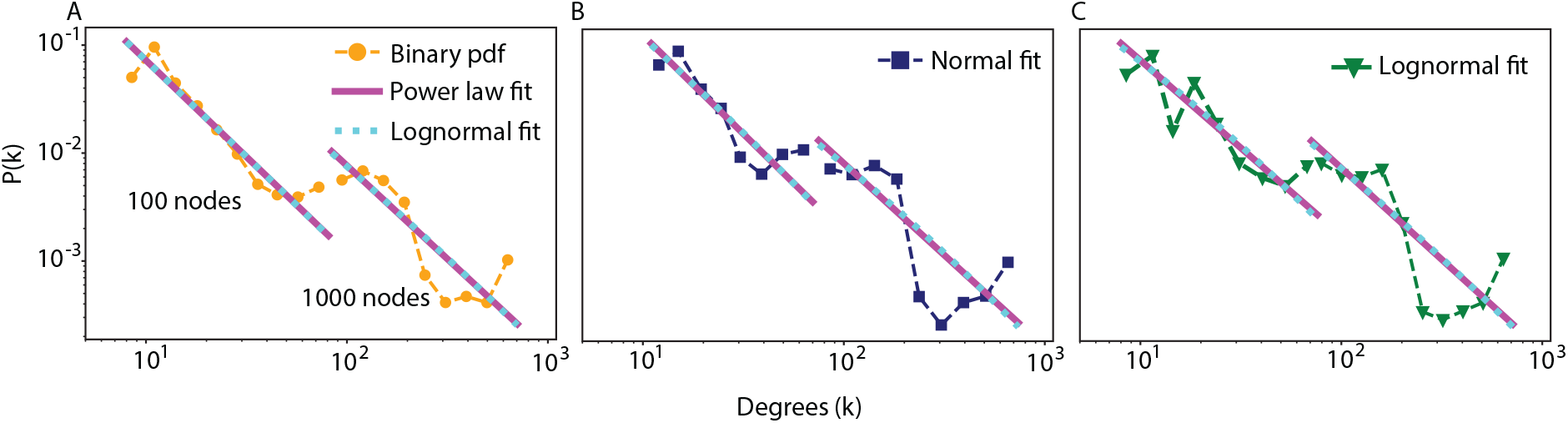
Power law and lognormal fits are better fits of the degree distribution of centralized networks compared to the exponential fit. Log-log plot of the degree distribution of an example 100 node network and a 1000 node network along with their power-law and lognormal fits for (A) binary, (B) normal and (C) lognormal networks.

**Fig S3.**
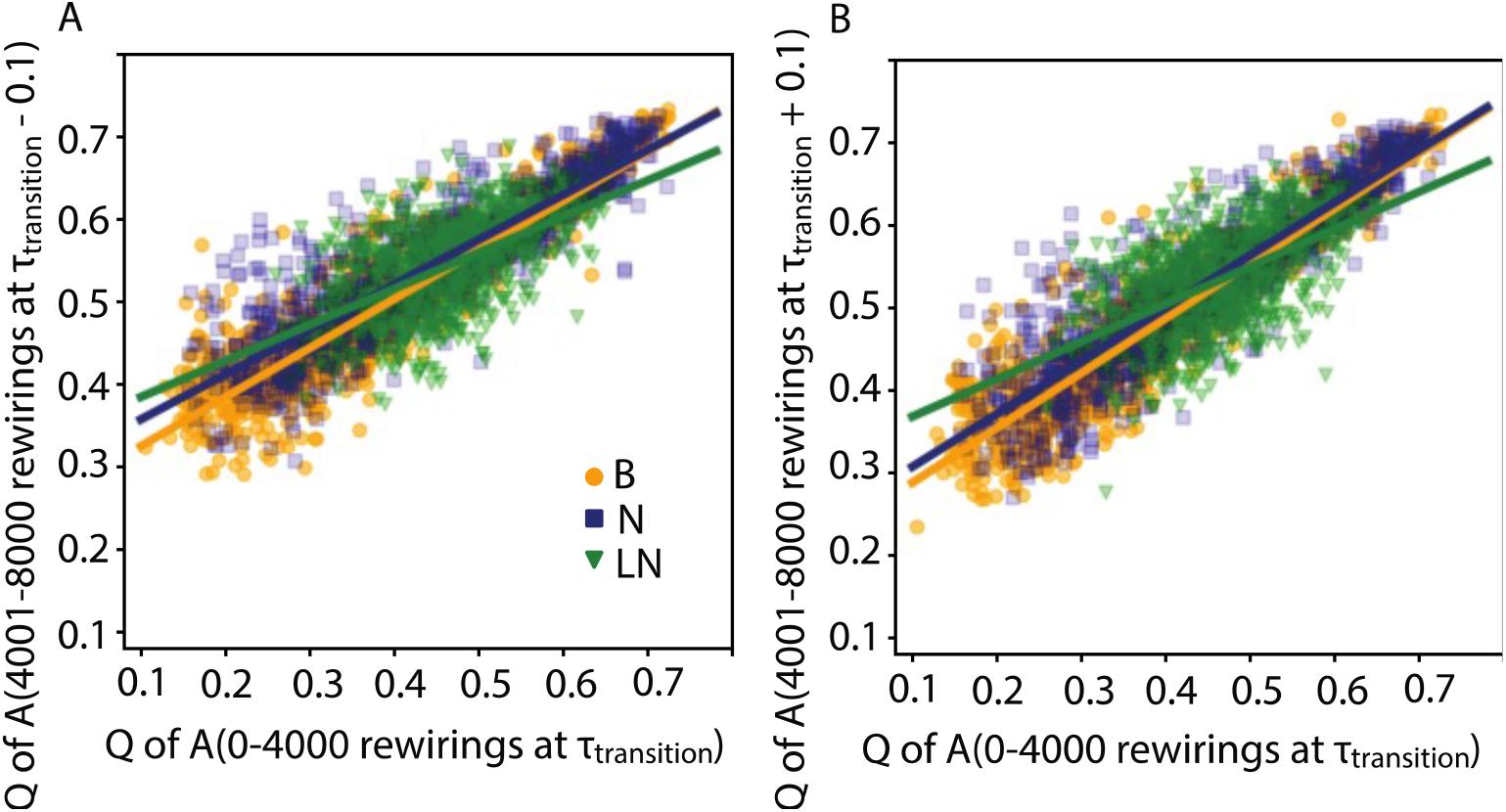
τ values near τ_transition_ do not affect the modularity index correlations seen at τ_transition_. A. Scatter plot showing strong positive correlation of the modularity indices between the same network after 4000 rewirings at τ_transition_ and an additional 4000 rewirings at τ_transition_-0.1 (towards the more modular direction) B. Same as A but for τ_transition_+0.1 (towards the more centralized direction). The lines in all cases indicate the best linear fits of the data.

**Fig S4.**
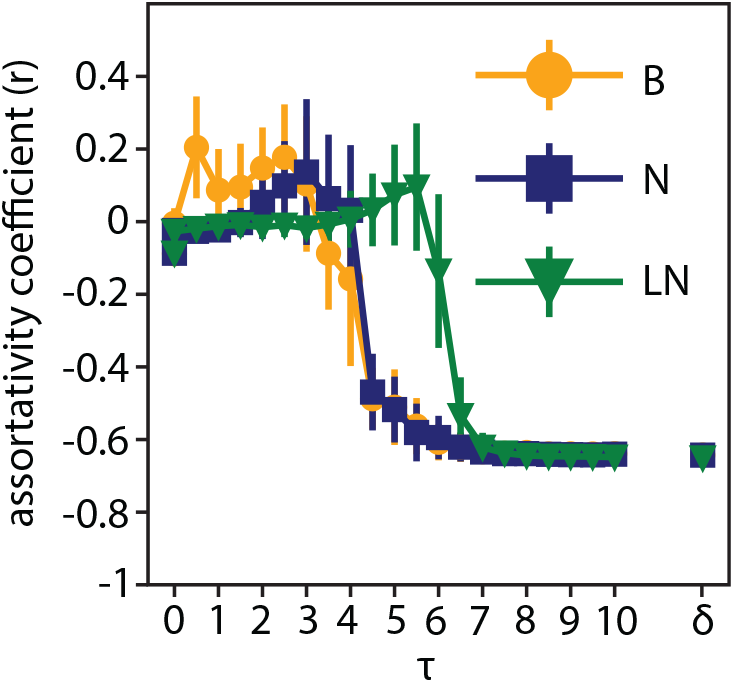
Networks are in two assortative states: a low and a disassortative one. The assortativity coefficient as a function of τ for binary, normal and lognormal networks. p_random_ = 0.2 for all networks.

**Fig S5.**
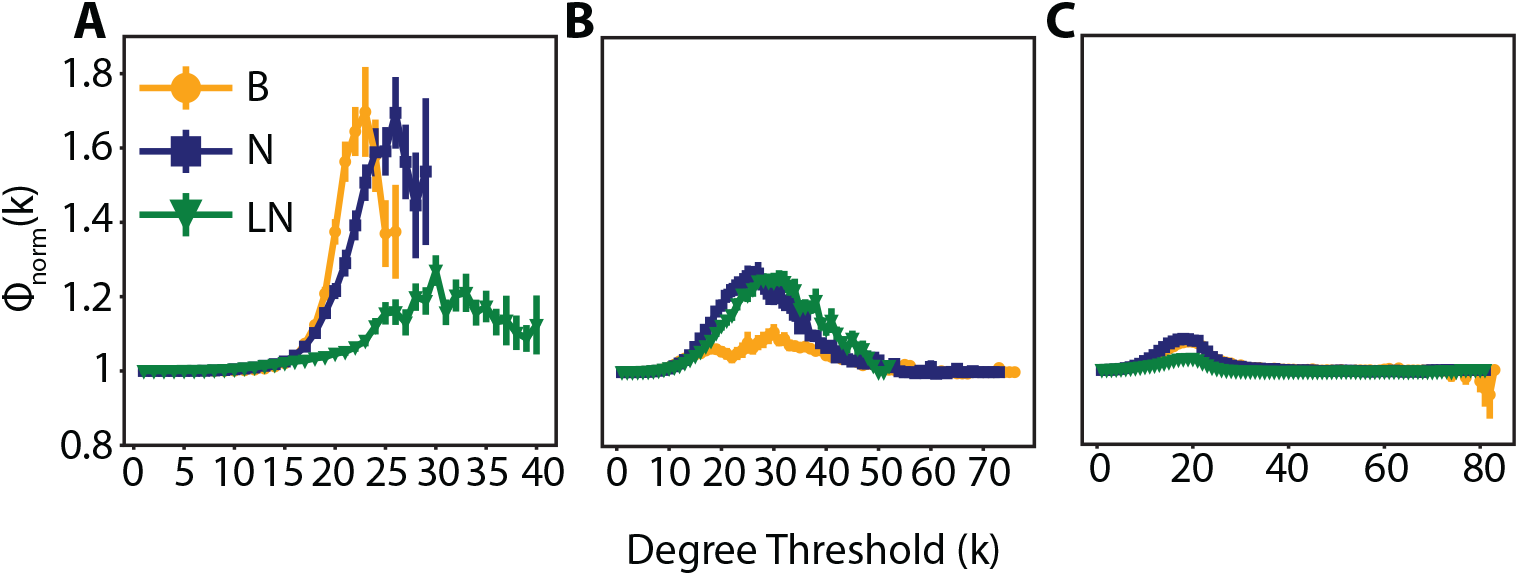
The weighted networks are rich club at the modular and transition state, the binary only at the modular state. Topological normalized rich club, Φ_norm_(k), for binary, normal and lognormal networks in the (A) modular (τ_binary_ = 2, τ_normal_ = 3, τ_lognormal_ = 4.5) (B) critical (τ_binary_ = 4.1, τ_normal_ = 4.15, τ_lognormal_ = 5.5) and (C) centralized (τ_binary_ = 5, τ_normal_ = 5, τ_lognormal_ = 7) state. In all cases p_random_ = 0

